# Subset-Based Analysis using Gene-Environment Interactions for Discovery of Genetic Associations across Multiple Studies or Phenotypes

**DOI:** 10.1101/326777

**Authors:** Youfei Yu, Lu Xia, Seunggeun Lee, Xiang Zhou, Heather M Stringham, Michael Boehnke, Bhramar Mukherjee

## Abstract

**Objectives:** Classical methods for combining summary data from genome-wide association studies (GWAS) only use marginal genetic effects and power can be compromised in the presence of heterogeneity. We aim to enhance the discovery of novel associated loci in the presence of heterogeneity of genetic effects in sub-groups defined by an environmental factor.

**Methods:** We present a p-value Assisted Subset Testing for Associations (pASTA) framework that generalizes the previously proposed *as*sociation analysis based on sub*set*s (ASSET) method by incorporating gene-environment (G-E) interactions into the testing procedure. We conduct simulation studies and provide two data examples.

**Results:** Simulation studies show that our proposal is more powerful than methods based on marginal associations in the presence of G-E interactions and maintains comparable power even in their absence. Both data examples demonstrate that our method can increase power to detect overall genetic associations and identify novel studies/phenotypes that contribute to the association.

**Conclusions:** Our proposed method can be a useful screening tool to identify candidate single nucleotide polymorphisms (SNPs) that are potentially associated with the trait(s) of interest for further validation. It also allows researchers to determine the most probable subset of traits that exhibit genetic associations in addition to the enhancement of power.

## INTRODUCTION

Genome-wide association studies (GWAS) are popular epidemiologic tools for studying the genetic architecture underlying a phenotypic trait [1]. Meta-analysis is a commonly used approach to combine genetic associations across multiple independent studies [2]. Fixed effect meta-analysis based on summary statistics is known to retain full efficiency of an analysis based on individual level data [3]. More recently, methods for aggregating association signals across multiple related phenotypes have also become more popular [4,5].

The *as*sociation analysis based on sub*set*s (ASSET) test, proposed by Bhattacharjee et al. [6], uses summary statistics from individual association analysis to develop a powerful test that allows the existence of both a subset of null results and effects in opposite directions in different individual tests. This approach essentially explores all possible non-empty subsets of available studies/traits and searches for the one that yields the strongest evidence of association while adjusting for multiple testing during such exhaustive search. One appealing feature of ASSET is that the most probable subset of studies/traits that exhibit genetic association can be identified in addition to the enhancement of power of detecting association signals. This method has been widely used in analyses of associations across multiple studies and phenotypes [7,8].

Exploiting gene-environment (G-E) interaction may help identify genetic variants that do not demonstrate very strong marginal effects due to environmental heterogeneity but may have stronger genetic effects in sub-groups defined by certain levels of an exposure [9–12]. Like most commonly used methods that screen only for marginal associations, ASSET does not account for potential G-E interactions in the testing procedure. If a genetic variant only affects a subgroup, for example the exposed group, then in the presence of such pure G-E interaction, the power of ASSET can be largely compromised. Several existing versions of 2-degree-of-freedom (2-df) tests for discovering genetic associations take G-E interactions into account. Kraft et al. [11] proposed a likelihood ratio test that jointly tests for genetic main effects and G-E interactions in case-control studies. Dai et al. [10] proposed a slightly different approach that is based on the sum of two Wald chi-square statistics for detecting marginal effects combined with G-E interactions. The latter approach has the advantage of incorporating G-E independence into the test statistic for G-E interaction in case-control studies to achieve increased power [13,14]. The literature on meta-analysis of G-E interactions or joint tests remains relatively limited [9–17], and none of these approaches attempt to specifically identify a subset of studies/traits that are most likely to have genetic associations and/or interactions. Current literature shows that the 2-df tests mentioned above can offer enhanced power in the presence of G-E interactions and do not lead to significant loss of power even when the interactions are absent [15,17].

In this paper, we propose a modification to ASSET that incorporates G-E interactions to increase the power of detecting genetic associations in combining summary data across studies of a given phenotype (say type 2 diabetes) or across multiple related phenotypes (say a set of lipid traits) measured in different studies. Our approach is similar to ASSET in terms of searching over all possible combinations of studies/phenotypes and thus inherits its advantage of identifying the maximally associated subset along with increased power for detecting the overall associations. The proposed framework uses p-values from the underlying tests as input instead of *z*-statistics that are used by ASSET, and is therefore referred to as p-value Assisted Subset Testing for Associations (pASTA). To be able to incorporate G-E interactions, pASTA uses 2-df tests as the underlying test statistics to be combined. In fact, the p-values can result from any statistical tests, including tests with multiple degrees of freedom. On the other hand, by using p-values, one loses the ability to incorporate the signs of coefficients (for both genetic main effect and G-E interaction) and the method is intrinsically two-sided. Since the key input variables are the p-values from underlying tests, pASTA can be used under any commonly used epidemiologic study design and we present our simulation studies and data examples for both case-control and cohort studies. Like ASSET, the analytical method in pASTA can handle a set of correlated p-values, enabling its validity in studies with overlapping subjects or multiple phenotypes measured on the same set of subjects.

The rest of the paper is organized as follows. We first describe the construction of our test statistic and derivation of the associated p-value. We present extensive simulation studies to compare the performance of our proposed method to ASSET and Fisher’s combined p-value approaches [18] with and without G-E interactions under various parameter settings. Specifically, we examine the type I error rates and the power to detect (1) the truly associated variant and (2) the truly associated subset of studies/traits for the given variant. We illustrate our proposed method with two data applications. The first is a meta-analysis of six case-control studies of type 2 diabetes (T2D) among European population, with 4,422 cases and 5,202 controls. We focus on two single nucleotide polymorphisms (SNPs) in the *FTO* gene (MIM 610966) related to obesity and their interactions with body mass index (BMI). The second application is a multiple-phenotype analysis of nine lipid-related quantitative traits using data of 5,123 individuals from the North Finland Birth Cohort 1966 (NFBC1966) Study. We investigate the association between two SNPs discovered by GWAS, one near the *LPL* gene (MIM 609708) and the other in the *APOB* gene (MIM 107730), and the nine lipid traits while taking the SNP-BMI interactions into consideration. We conclude the paper with a discussion.

## MATERIAL AND METHODS

### Notation

We consider *K* studies, each with sample size *n*_*k*_. We allow the studies to have a set of overlapping subjects. Let *Y*_*ki*_, *G*_*ki*_ and *E*_*ki*_ denote the underlying phenotype, the given genetic variant and the environmental factor, respectively, measured for the *i*th subject in the *k* th study. The trait(s) *Y* can either be disease status in a case-control study or binary or quantitative trait(s) measured in a cohort study. The genetic variant *G* may be coded as binary (under a dominant or recessive susceptibility model) or as allele count (under the additive model), and for the latter case, we treat *G* as continuous dosage. The environmental factor *E* can either be categorical or continuous. We mainly consider two models for each of the *K* studies: the marginal genetic association model

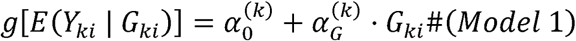

and the joint model with G-E interaction

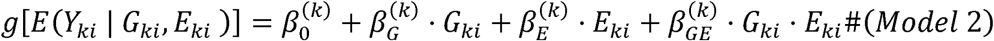

where *i =* 1,*…,n*_*k*_ and *k =* 1,*…, K*. The choice of the link function *g*(*·*) depends upon the type of *Y*. Adjusted covariates are dropped from the presentation for simplicity of notations. Note that the *K* studies in this setup can be replaced by *K* traits in a single study of size *n*, and the inferential procedure for each of the *K* traits stays the same with *n*_*k*_ *= n* for each *k = 1,*…, *K*.

We primarily consider three types of association tests for each of the *K* studies: 1) marginal genetic effect in *Model 1* (MA), with null hypothesis (*H*_0*k*_) being 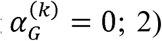 genetic main effect and G-E interaction in *Model 2* (JOINT) [11], with *H*_0*k*_ being 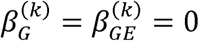 and 3) marginal genetic effect in *Model 1* and G-E interaction in *Model 2* (MA+GE) [10], with *H*_0*k*_ being 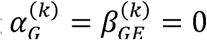 MA is a 1-df test while JOINT and MA+GE are 2-df tests. A summary description of the different tests and their corresponding null hypotheses, test statistics and distributions under the null are presented in Supplemental Table S1. The test of the form MA+GE is typically the sum of two Wald chi-square test statistics that are known to be asymptotically independent and can thus be combined to yield a 2-df chi-square statistic [19].

For a case-control study, the coefficient of interaction 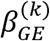 can be estimated using case-control (CC), case-only (CO) [13], or empirical Bayes (EB) [14] estimators, respectively denoted as 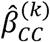, 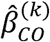 and 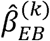. It has been proved that 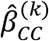 and 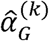 are asymptotically independent, and so are 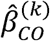 and 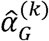 when we assume G-E independence and a rare disease [19]. As such, for case-control studies we use 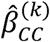 and 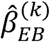 separately as two possible choices for the estimator of 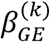 when computing the test statistic of MA+GE, and denote these two types of tests as MA+CC and MA+EB, respectively [10]. Note that the use of G-E independence requires the use of a retrospective likelihood of the form (*G,E*)*|D*, which is not valid unless the sampling is conditional on *D*. A cohort analysis is always based on *D*|(*G, E*) and we cannot model the stochastic distribution of (*G, E*). Therefore, for a cohort study with multiple quantitative or binary phenotypes, the use of G-E independence does not lead to enhanced power and as such the EB estimator does not apply.

### p-value Assisted Subset Testing for Associations (pASTA)

Regardless of the test statistic chosen for each study, the underlying analytical framework of pASTA starts with a set of p-values: *P*_*k*_, *k = 1,…, K*. In our setting they are obtained from the three tests, MA, JOINT, or MA+GE. For pASTA, we define the *z*-statistic for the *k* th study as

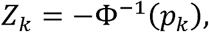

where *Φ*(*·*) is the cumulative distribution function (CDF) corresponding to the standard normal distribution. That is, smaller p-values indicate larger *z*-statistics, and the null hypothesis *H*_0*k*_ should be rejected if *z*_*k*_ is large in the positive direction. For any non-empty subset *S* ⊆ {1,*…, K*}, let

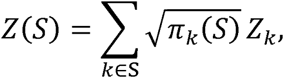

where we consider 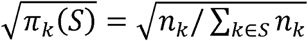 as the weight of *z*_*k.*_ Other options for the weights 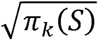 can also be accommodated in this framework. For example, when cases and controls are not of equal numbers, *n*_*k*_ can be substituted by the effective sample size [2]. The test statistic for evaluating the overall association of a given genetic variant is defined as

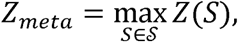

and is obtained by searching over all possible *| 𝒮 | = 2*^*K*^ –1 non-empty subsets of studies.

It is challenging to characterize the exact distribution of *z*_*meta*_ in a closed form, especially when the number of studies *K* is large. Therefore, we borrow the idea followed in ASSET and use the discrete local maxima (DLM) procedure [20] to obtain a conservative upper bound for the meta-analytic combined p-value. The DLM procedure automatically accounts for the multiple comparison issue. The detailed derivation of the combined p-value based on *Z*_*meta*_ is included in Appendix. In brief, let *γ ∈ Γ* be a *K*-dimensional vector, each of whose components takes value in {*0, 1*}, and let *S*_*γ*_ be the subset of studies whose corresponding coordinates in *γ* are 1. A neighbor of *S*_*γ*_ is defined as a subset in *𝒮* that can be obtained by adding or dropping one study to or from *S*_*γ*_. Then *z*(*S*_*γ*_*) is called a local maxima if it is greater than all *z*(*S*_*γ*_)’s where *S*_*γ*_ refers to any possible neighbor of *S*_*γ*_*. For an observed test statistic *z*_*meta*_ *= T*_*obs*_, the DLM approximated p-value can be expressed as (see Equation A1 in Appendix)

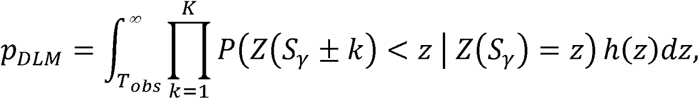

where, *h*(·) is the probability distribution function of *Z*(*S*_*γ*_). This DLM-based p-value is actually an upper bound to the exact one (Equation A1 in Appendix). Hence the overall procedure is conservative in terms of type I error.

Each of the terms of the form *P*(Z(*S*_*γ*_± *k*) < *z* | Z(*S*_*γ*_ *= z*) can be evaluated based on the conditional normal distribution of *Z*_*k*_ given Z(*S*_*γ*_) (Equations A2 and A3 in Appendix). Thus all we need to evaluate is the joint bivariate distributions of *Z*_*k*_ and Z(S_γ_) for *k =* 1,*…, K*. Note that *Z*_*k*_ interaction) in study *k*. The set of *Z*_*k*_’s are independent if the studies are independent. For multiple follows the standard normal distribution under the null hypothesis that there is no association (and studies with overlapping subjects or multiple phenotypes measured on the same set of individuals, *Z*_*k*_’s will be correlated with variance one and correlation matrix ∑ with entries say ρ_*kk’*_ *= Corr*(*Z*_*k*_, *Z*_*k’*_).

Let ***L =***(*L*_1_,…, *L*_*k*_), where *L*_*i*_ =1 if the *i*-th study is in *S*_*γ*_ and *L*_*i*_ = 0 otherwise. Let ***Z*** = (*Z*_1_,…, *Z*_*K*_) follow a multivariate normal distribution *MVN*_*K*_ (**0**, ∑). In particular, if all studies/traits are independent, then ∑ is a *K*×;*K* identity matrix. Let ***w*** = (w_*1*_,…, w_*K*_) be the vector of weights for the element in ***L***. 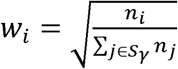 if *L*_*i*_ = 1 and 0 if *L*_i_ = 0. Let ***e***_*k*_ be a vector of length *K* where the *k*th element equals 1 and 0 elsewhere, e.g. ***e*** _*2*_ = (0,1,0,.., 0)^*T*^. Let 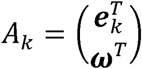,which is a *2 ×; K* matrix. Under the null hypothesis,

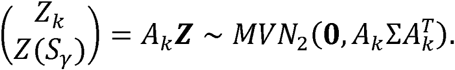

Several methods have been proposed to estimate the correlation matrix ∑. Lin and Sullivan ([21] provided a formula to estimate the correlation between a pair of studies using the information of shared cases and controls when the number of overlapping subjects is known. They also discussed a modified version of the formula for quantitative traits in a cross-sectional study. Bhattacharjee et al. [6] extended Lin and Sullivan’s formula for case-control studies to accommodate the situation where cases of one study serves as controls of another. However, these estimates will likely be inaccurate when the overlap fraction is large, and therefore do not directly apply to multiple-phenotype analysis, where the phenotypes are measured on the same set of subjects and the overlap fraction is 100%. In this case, associations and thus large positive,-statistics, an intuitive way to approximate the correlations of since large positive values of phenotypic measurements usually indicate strong positive genetic *Z*_*k*_’s is to use the phenotypic correlations [22], as is done in ASSET. We adopt this approach for our simulation studies and the second data example to estimate the correlation matrix ∑ corresponding to ***Z***. In other words, if 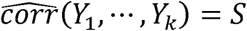, then we let 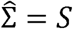.

### Comparison of pASTA and ASSET

We would like to point out a few features of pASTA as compared to ASSET when one is only interested in marginal genetic association estimated by *Model 1*. First, ASSET and pASTA use the *Z*-We would like to point out a few features of pASTA as compared to ASSET when one is only statistics in different ways (Table S2). The *Z*-statistics used in pASTA no longer differentiate between possible directions of association, while the signs of the *Z*-statistics used in ASSET are consistent with those of. 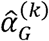’s. ASSET can detect genetic associations in either one direction (one-sided) or both directions (two-sided). For the latter case, ASSET searches for the subsets of studies that yield the strongest evidence separately in both directions. It obtains a p-value for each direction, and combines the two p-values using Fisher’s combined p-value method [18]. pASTA does not have this feature. A more detailed comparison of the *Z*-statistics, test statistics and analytic expression for the DLM-based p-values of the two methods are presented in Table S2. The extension in this paper is mainly to incorporate interaction while testing association using 2-df tests in the ASSET framework via the use of p-values.

### Simulation studies

#### Simulation Study 1: Combining Signals Across Multiple Independent Case-Control Studies

##### Methods Considered

We first assess the performance of the proposed pASTA approach for combining summary results from multiple independent case-control studies that have no overlapping subjects or relatedness. We compare pASTA to three classes of alternatives (Table 1): ASSET, Fisher’s combined p-value method [18], and the gold standard test which assumes that the true subset of non-null studies and the true model from which the data are generated are known a priori, to reflect the maximum achievable power for benchmarking each method. Given, independent p-values *p*_*1*_,*…, p*_*k*_, the chi-square test statistic of Fisher’s method follows a 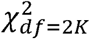 distribution under the global null. Bhattacharjee et al. [6] has already compared inverse-variance weighted meta-analysis with ASSET and Fisher’s method, showing that the former is not as powerful as the latter two. Therefore, we do not include standard fixed effect inverse-variance weighted meta-analysis method in our comparative study. We use several choices of 2 df tests as described in Table 2 and expanded in Table S1.

**Table 1.**
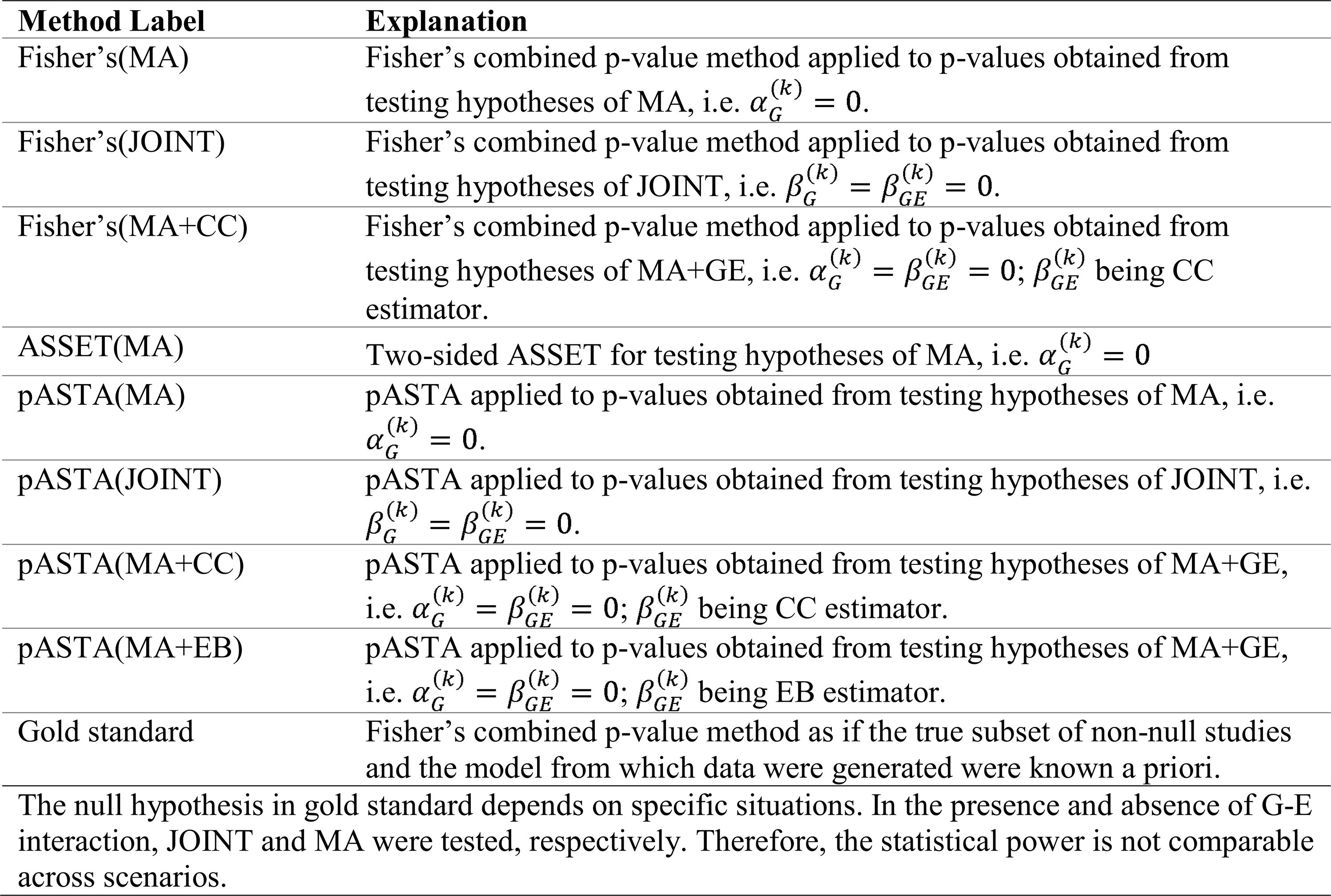
Methods Considered in Simulation Studies

**Table 2.**
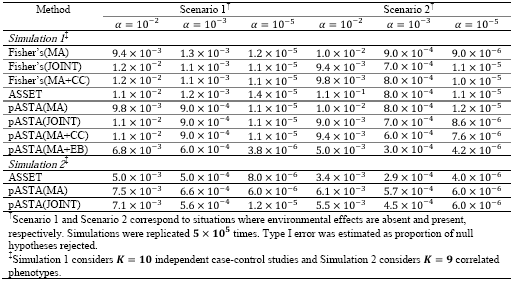
Type I Error Rates at Various Significance Levels for the Two Simulation Studies

##### Data Generative Model

We consider *K* =10 independent studies with 6,000 cases and 6,000 controls. The disease status *D*, the genetic marker *G* and the environmental factor *E* are assumed binary (1 for presence and 0 for absence) for simplicity. Following the standard approach of simulating case-control data with G-E interaction [9], we generate the data from a logistic regression model

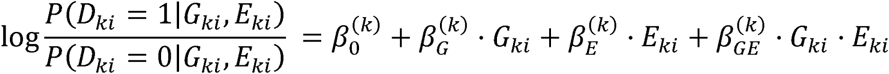

for *k* = 1,…, *K* In addition, we let θ_*GE*_, *p*_*G*_ and *p*_*E*_ denote the G-E association, the frequency of *G* = 1(this probability depends on the frequency of the risk allele, we only consider a dominant model in the simulation study) and the prevalence of environmental factor (*E* = 1) in controls, respectively. We specify the parameters *θ*_*GE*_, *p*_*G*_, *p*_*E*_, 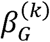, 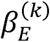 and 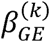, and determine the odds ratios of marginal genetic association (OR_G_) and environmental association (OR_E_) using the formula in Boonstra et al. [9]. In all of our simulation studies, we assume G-E independence among the control population, i.e., *θ* _*GE*_ = 0.

##### Evaluation of Type I Error

To assess the type I error rate of each method in the presence or absence of environmental effects, we generate data under the null hypothesis of no genetic effects, namely, set 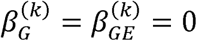 in *Model 2*. We consider two scenarios (Table S3), one with no environmental factor and hence no G-E interaction and the other with OR_*E*_ = 1.4. All ten studies share the same parameters in each scenario (Table S3). We evaluate the type I errors at three levels of significance:.*α* = 10^*-2*^ and 10^*-3*^ resembling a candidate gene study, and 10^*-5*^resembling large scale exploration. We simulate 5×10^*5*^replicated datasets for each scenario.

##### Metrics for Power Analysis

We compare the power of pASTA with the other methods displayed in Table 1 from two perspectives: the power of detecting overall genetic associations and the accuracy of identifying the exact subset of non-null studies. Here the non-null studies refer to studies in which 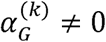 (and hence 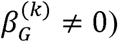) or 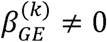. The specific definitions of the two metrics are given in Appendix. We also report the sensitivity and specificity of identifying the subset of non-null studies, defined as the proportion of non-null studies that are correctly identified by subset-based approaches, and the proportion of truly null studies that are declared null, respectively.

##### Power for Detecting Marginal Association

Prior to consideration of G-E interaction we extensively investigate the relative performance of ASSET and pASTA(MA) in detecting marginal genetic associations in a case-control setting where the data are generated under *Model 1*. We consider two scenarios in this simulation, each with *K* = 10 independent studies and *k*_*o*_ = 4 of them having true associations. The prevalence of the genetic variant *p*_*G*_ is 0.2. In the first scenario, we assume that the reverse the sign of the log-odds ratio parameter of genetic effect (*β*_*G*_) in two of the four studies. In both marginal genetic effects in the four non-null studies are all in the positive direction. In the second, we scenarios, the absolute values of log odds ratio of marginal associations 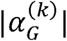 are the same for all non-null studies, ranging from log1.05 to log1.2, and 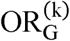 is incremented by 0.01. Since in reality it is possible that all studies have non-null signals, we also carry out a simulation study where *k*_0_ = 10. We use *α* = 0.001 as the significance level, which resembles a candidate gene study with 50 SNPs, and repeat the simulation 2,000 times for each parameter configuration.

##### Power for Detecting Association After Incorporating Interaction

We then consider both, and, in our model and the whole set of 1-df and 2-df tests as listed in Table 1. We consider five (Table S4) to evaluate the power of pASTA. Scenario 1 assumes no G-E interaction and a moderate scenarios marginal genetic effect (OR_*G*_). In Scenario 2, we set parameters to induce large G-E interaction but small marginal genetic effect. Scenario 3 considers the same magnitude of interaction as Scenario 2, but with zero genetic and environmental main effects, which result in a smaller marginal genetic effect compared to Scenario 2. Scenario 4 assumes strong G-E interaction and protective genetic main effect, the two of which cancel out and result in an odds ratio of marginal genetic effect close to one. Scenario 5 considers negative interaction and positive genetic main effect, which lead to a nearly-null marginal genetic effect. The last two scenarios represent less common case-control settings, where the genetic associations exist in opposite directions for exposed and unexposed groups. The number of non-null studies ranges from two to seven. Simulations are repeated 2,000 times for each scenario, and significance level is *α* = 0.001.

#### Simulation Study 2: A Multiple-Phenotype Analysis in a Cohort Study

##### Methods Considered

We compare the performance of pASTA and ASSET in a cohort study with a set of multiple correlated traits. In this case, the test statistics of Fisher’s method no longer follows a 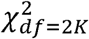 distribution under the null hypotheses due to the correlations among *p*_*1*_,*…, p*_*k*_.. Resampling-based approaches can be used to obtain the null distribution when correlations exist [23], but in this situation we are dealing with significance level at the magnitude of 10^*-S*^ or smaller. It is computationally intensive to obtain a null distribution to evaluate the p-value at such a stringent threshold. Therefore, we exclude comparison with Fisher’s method from Simulation Study 2.

##### Data Generative Model

We consider a multiple-phenotype study with 6,000 subjects and *K* = 9 correlated traits. The outcome *Y* is generated from a multivariate linear regression model

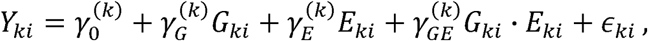

where (∈_1*i*_,…, ∈_*ki*_)^T^ ∼*MVN*(0,Ω). We use the empirical phenotypic correlations from the NFBC1966 study (Table S5) as the generative covariance matrix ?.

##### Type I Error and Power Analysis

Similar to Simulation Study 1, the type I error rates are estimated in the presence and absence of environmental factor (Table S6). The power of these subset-based approaches are evaluated in two scenarios (Table S7) using the same metrics as used in Simulation Study 1. In Scenario 1, no interaction is involved, while in Scenario 2, interaction comes into play.

### Data Analyses

We illustrate our methods by using data from two genetic association studies. The first is a meta-analysis of six independent case-control studies of T2D and the other considers a multiple-phenotype analysis of nine quantitative lipid-related traits measured on a population-based cohort. In both examples, we compare the performance of pASTA with other competing methods as described in Table 1.

### Application to Case-Control Studies of Type 2 Diabetes

This analysis includes data from a subset of subjects in six independent studies of T2D: FIN-D2D 2007 (D2D2007), The DIAbetes GENetic Study (DIAGEN), the Finland-United States Investigation of NIDDM Genetics Stage 2 (FUSION S2), The Nord-Trøndelag Health Study 2 (HUNT), the METabolic Syndrome In Men Study (METSIM) and the Tromsø Study (TROMSO) [24]. We investigated the associations of T2D and two SNPs in the *FTO* gene, rs6499640 and rs1121980, respectively after taking into account potential interaction effects with BMI. These two SNPs were chosen as candidates because they showed significant interactions with BMI in a previous study [25]. Descriptive statistics of the six studies are summarized in Supplemental Table S8. In particular, the total sample size of the six studies was 9,624, with 4,422 cases and 5,202 controls. Sample sizes of the six studies vary from 1,058 to 2,219 and the case to control ratios vary from 0.37 to 1.75. The minor allele frequencies (MAF) of rs6499640 and rs1121980 in controls range from 0.37 to 0.43 and 0.39 to 0.48, respectively, across the studies.

The outcome of interest is the presence, 1 or absence, 0 of T2D. Each SNP is coded as *G =* 0,1,2, representing the number of minor alleles in rs6499640 or rs1121980 for each subject assuming an additive genetic susceptibility model. Let *E* denote BMI, *A* denote age in years, and *S* represent an indicator of sex (Male = 1). For the *i*-th individual in the *k*-th study, the standard logistic regression model incorporating G-E interaction is specified as

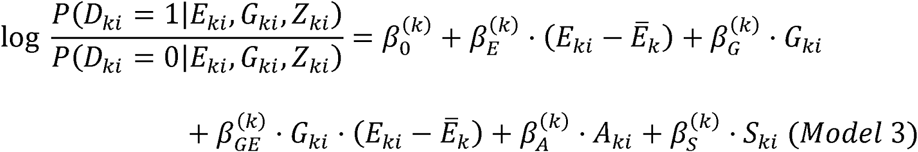

and the standard logistic regression model of marginal SNP effect (adjusted for BMI, age and gender) is specified as

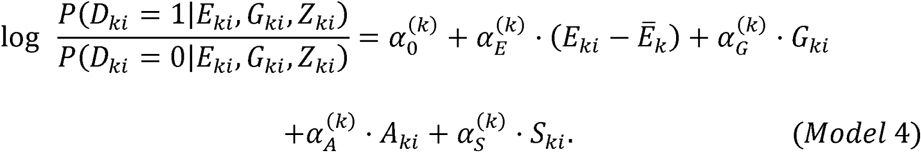

*Model 3* and *Model 4* were fitted separately to the six studies. Then the various meta-analysis methods were applied to the summary statistics. Note that the METSIM study only contains male subjects, so the sex indicator was removed from the two models for this study.

### Application to the North Finland Birth Cohort 1966 Study

We obtained the genotype and phenotype data of the NFBC1966 study from dbGaP with accession number phs000276.v2.p1. The NFBC1966 data contain 5,402 individuals and 364,590 SNPs, with multiple metabolic traits measured on subsets of individuals. Among these traits, we considered nine quantitative traits in the present study as phenotypes: C-reactive protein (CRP), glucose, insulin, total cholesterol (TC), high density lipoprotein (HDL), low density lipoprotein (LDL), triglycerides (TG), systolic blood pressure (SBP), and diastolic blood pressure (DBP). Body mass index (BMI) was the environmental factor of interest. Following previous studies [26,27] we excluded individuals with missing phenotype data or having discrepancies between reported sex and sex determined from the X chromosome. We excluded SNPs with a minor allele frequency less than 1%, having missing values in more than 1% of the individuals, or with a Hardy-Weinberg equilibrium p-value below 10^*-4*^. This left us with 5,123 individuals and 319,147 SNPs. For each phenotype in turn, we quantile transformed the phenotypic values to a standard normal distribution, regressed out sex, oral contraceptives and pregnancy status effects [26], and quantile transformed the residuals to a standard normal distribution again. We replaced the missing genotypes for a given SNP with its mean genotype value. To account for potential relatedness in the Finnish population one can use a kinship matrix in a linear mixed model to account for population admixture. As an alternative, we find that using linear regression models with principal components (PCs) as covariates also effectively controls for population stratification in the data. For example, with ten PCs, the genomic control factors for the ten phenotypes ranged from 0.96 to 1.01, with a median value of 0.99. Therefore, we extracted the top ten PCs from the genotype matrix as covariates to control for potential population stratifications.

The linear regression model with G-E interaction is specified as

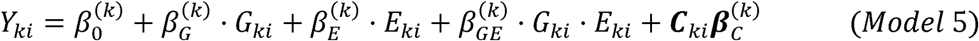

and the linear regression model of marginal SNP effect (adjusted for BMI and PCs) is specified as

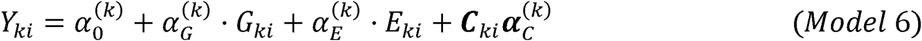

where *k* indexes phenotypes, *i* indexes subjects, *E* denotes BMI, *G* denotes the number of minor alleles, and ***C*** denotes the vector of top ten PCs. We performed a genome-wide analysis of multiple phenotypes by fitting *Model 5* and *Model 6* separately to each of the nine phenotypes for all 319,147 SNPs in the data. Then for each SNP we calculated the meta-analytic p-values using ASSET and pASTA, respectively. Correlations among the resultant *Z*-statistics of the nine phenotypes were estimated using observed phenotypic correlations.

After the genome-wide screening using the nine phenotypes separately, we performed another genome-wide association analysis by analyzing all ten traits jointly with the multivariate linear mixed model implemented in GEMMA [28,29] and compared the results of the two GWAS. We then selected the top 100 SNPs that showed the strongest associations for subsequent interaction analysis. We performed SNP-BMI interaction analysis on these 100 SNPs.

## Results

### Simulation Study 1: Combining Signals Across Multiple Independent Case-Control Studies

#### Type I Error

In both scenarios, empirical type I error rates at thresholds. *α* = 10^*-2,*^ 10^*-3,*^ and 10^*-S*^ are well-preserved for all methods except pASTA(MA+EB) (Table 2), which tends to be conservative (e.g. type I error rate being 3.8 ×; 10^*-6*^ at the level of 10^*-5*^). This has been noted in previous studies of EB-type adaptive methods [14].

#### Power Comparisons of ASSET and pASTA Based on 1-df Tests for Marginal Genetic Association

We extensively investigate how pASTA(MA) performs compared to ASSET when no environmental factors are considered (i.e. data generated under *Model 1*). The power of pASTA to detect overall genetic associations is almost the same as that of ASSET as the odds ratio of the marginal genetic effects varies, regardless of the directions of the genetic effects (Figures S1A, S1E). However, there is an observable difference in their accuracy of determining the exact subset of non-null studies for both scenarios (Figures S1B, S1F). When the marginal genetic effects of non-null studies exist in the same direction (in this case positive direction), the accuracy of pASTA to determine the correct subset increases as the signal grows stronger, and reaches an empirical power of 0.72 as the odds ratio goes up to 1.2. However, two-sided ASSET is rarely able to identify the exact true subset, and the corresponding power stays around zero as the odds ratio varies. The main reason for such low accuracy is that two-sided ASSET searches for associated subset in both positive and negative directions, identifies a subset that gives the largest |*Z*(*S*)| for each direction, and takes the union of the two subsets as the final identified subset regardless of the significance of association in either direction. As a result, two-sided ASSET generally identifies more false positive studies when the significant associations are all in one direction. For example, when associations only exist in the positive direction in a subset of studies, the log odds ratios in the truly null studies are likely to be estimated as negative, and thus some null studies will be selected by two-sided ASSET. When the associations exist in opposite directions, two-sided ASSET has improved power to identify the correct subset of studies, but pASTA still consistently outperforms ASSET (Figures S1B, S1F). For both scenarios, the sensitivity of ASSET consistently stays higher than that of pASTA, while the specificity of ASSET is relatively low (Figures S1C, S1D, S1G, S1H). These results are intuitively reasonable, since sensitivity is the proportion of non-null studies that are correctly identified and does not require that the exact subsets to be identified.

When all of the ten studies are non-null (Figure S2A), pASTA is as powerful as ASSET in detecting signals of marginal associations, and only slightly less powerful than Fisher’s method when the odds ratio of marginal genetic effects is small (<1.1).

#### Power Comparison of All Methods with Data Generated under Model 2

##### Detection of Association

We consider five scenarios (Table S4) to evaluate the power of detecting overall genetic associations. When there exists only marginal genetic effect but no G-E interaction (Scenario 1), 1-df methods are slightly more powerful in discovering the overall associations than 2-df methods when the number of non-null studies *k*_0_ *≤* 5. For example, pASTA(JOINT) reaches a power of 0.85 when four out of the ten studies are non-null, which is 9.1%, 7.1%, and 9.9% lower than the power of Fisher’s(MA), ASSET, and pASTA(MA), respectively (Figure 1A). This loss of power results from a penalty of the additional degree of freedom incorporated in the testing procedure in the absence of G-E interaction. In Scenario 2, when G-E interaction exists and marginal genetic effect is moderate, all 2-df methods provide substantial power gain over 1-df methods (Figure 1B). For example, when *k*_0_ = 7, the power of pASTA(MA+EB) reaches 0.99, while that of ASSET stays around 0.6. In Scenario 3, where the marginal genetic effect is even smaller, the power of all methods is compromised, with pASTA(MA+EB) achieving the best power of 0.79 when *k*_0_ = 7. The gold standard test has the second best performance with the power being 0.61 when *k*_0_ = 7. All 1-df methods barely detect any signals (Figure 1C). When the marginal association is nearly null but sub-group effects of genetic factor present in opposite directions in the exposed and unexposed group (Scenario 4), 2-df methods are more powerful in detecting signals than their 1-df counterparts. Specifically, pASTA(MA+EB) performs the best when *k*_0_,3, already achieving a power of 0.82 when,_*O*_ 3. Other 2-df methods perform slightly worse, with their power ranging from 0.44 to 0.52 when,_*O*_ 3, but they still yield power greater than 0.8 when,_*O*_,5. All 1-df methods fail to detect the associations (Figure 1D). In a much less common situation where “cross-over” interaction exists (Scenario 5), the empirical power curves show a similar trend (Figure 1E) to those in Scenario 4. It is instructive to note that for Scenario 2-5 where the interaction effects are present, we tested the null hypothesis of JOINT for the gold standard test. Therefore, pASTA(MA+EB) outperforms the gold standard when *k*_0_ is large since EB estimator exploits the putative G-E independence and can be more efficient in this case.

##### Subset Identification

When the environmental factor is absent (Scenario 1), pASTA(MA) achieves the highest accuracy in identifying the exact subset of non-null studies, with the corresponding power being constantly around 0.4 (Figure 1F). The sensitivity of the subset-based methods, whether they are 1-df or 2-df, are close to one another as the number of non-null studies varies, and all of the methods are able to identify more than 80% of the non-null studies (Figure S3A). The specificity of ASSET is much smaller than that of the other subset-based methods based on pASTA (Figure S3F). When G-E interaction comes into play (Scenario 2-5), pASTA(MA+EB) performs the best in identifying the exact subset and has both the highest sensitivity and specificity across the four scenarios. In particular, in Scenario 4 where the genetic variant has opposite effects on exposed and unexposed groups, the highest empirical probability of pASTA(MA+EB) to select the correct non-null studies is 0.46 when three out of the ten studies are non-null. pASTA(JOINT) and pASTA(MA+CC) have the second-best performance in terms of determining the exact subset of non-null studies as well as sensitivity and specificity across Scenarios 2-5. It is noteworthy that in all five scenarios, the specificity of ASSET is the lowest among the subset-based methods, which indicates that ASSET tends to give more false positives of non-null studies than pASTA.

#### Simulation Study 2: A Multiple-Phenotype Analysis in a Cohort Study

##### Type I Error

The results of the type I error rates are presented in Table 2. In general, the type I error rates of all methods tend to be conservative at all thresholds under evaluation. The only exception is that pASTA(JOINT) yields a type I error of 1.2×10^*-5*^at the level of 10^*-5*^ when no environmental factor is considered. The conservativeness of the type I errors may be due to the inaccurate estimation of ∑, the correlation matrix of the *Z*-statistics.

#### Power Comparison

##### Detection of Association

Figure 2A reveals that in the absence of interaction, pASTA(MA) outperforms the other two methods when over five out of the nine traits have genetic associations. The power of ASSET drastically falls when the number of non-null traits exceeds five due to computational issues of the ASSET package. According to Figure 2E, when interaction exists, pASTA(JOINT) provides great power gain in detecting the overall associations and reaches a power of 0.92 when five out of the nine traits are non-null. The power of the 1-df methods stays below 0.9.

##### Subset Identification

Similar to what we observe in Simulation Study 1, the accuracy of ASSET in identifying the exact subset of non-null traits are constantly close to 0 in both scenarios (Figures 2B, 2F). In Figures 2C and 2G, the sensitivity of the pASTA approaches is close to one another and stays above 0.8 as the number of non-null traits varies, which indicates that these methods are able to identify over 80% of the non-null traits in one study. On the other hand, the sensitivity of ASSET decreases as the number of non-null traits increases. The curves of specificity (Figures 2D, 2H) show similar trends to those of sensitivity.

### Application 1: Type 2 Diabetes Data

The exponential of the parameter estimates (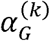, 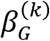 and 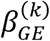) their corresponding 95% confidence interval (CI) and p-values obtained from *Model 3* and *Model 4* for each of the six studies are summarized in Supplemental Table S9. The EB estimates of the interaction terms and their associated Wald chi-square statistics were obtained using the R package CGEN [30]. For SNP rs6499640, the odds ratios of marginal genetic effect range from 0.89 to 1.11, and none of them are significant at the level of 0.05. When the interaction between SNP and BMI is included in the model, the odds ratios of genetic main effect do not change remarkably from those of marginal genetic effects. However, despite the small effect sizes of interactions in all six studies, the interaction effects are statistically significant in D2D2007 for CC method (p=0.017) and in FUSION S2 for CC (p=0.0006) and EB (p=0.003) methods. The SNP rs1121980 has significant marginal associations with T2D in FUSION S2 (*OR*_G_ = 1.19, p=0.011) and HUNT (*OR*_*G*_ 1.21, p=0.020), while none of the studies show significant SNP-BMI interactions for this SNP.

We then meta-analyzed the results from the six studies by applying all methods in Table 1, and we present the results of the eight methods in Table 3. For SNP rs6499640, all 1-df methods give insignificant p-values for marginal genetic associations with T2D at the level of 0.05. As for 2-df methods, pASTA(JOINT) gives the smallest p-value (p=0.008), and pASTA(MA+CC) give a slightly higher p-value (p=0.009). The result of pASTA(MA+EB) is not statistically significant (p=0.090), which may be due to the presence of G-E correlation in this example. All 2-df pASTA methods identify D2D2007 and FUSION S2 as the non-null studies, which is consistent with what we observe from the results of 2-df tests for each study separately. We also investigated the odds ratios of T2D across different levels of BMI by fitting a logistic regression adjusting for age, gender, and the studies to which the subjects belong. Supplemental Figure S4 shows that the marginal odds ratio for the SNP rs6499640 is close to 1, but when G-E interaction is taken into account, this SNP reveals a significant protective effect at the level of 0.05 when BMI equals 20 and 25. When BMI equals 30, the odds ratio of T2D becomes positive and is significantly different from the one when BMI equals 20.

**Table 3.**
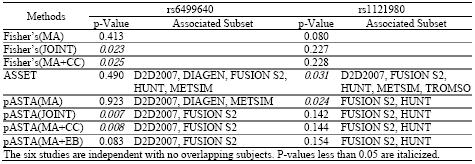
Meta-analysis of Six Case-control Studies of Type 2 Diabetes

When there is no evidence for G-E interactions in individual studies, as with SNP rs1121980, all 2-df methods fail to detect the overall genetic associations, but the p-values obtained from 2-df pASTA are smaller than those from 2-df Fisher’s p-value combined methods. Furthermore, the p-value given by pASTA(MA) (p=0.024) is smaller than those of Fisher’s(MA) (p=0.080) and ASSET (p=0.031). pASTA also provides more plausible subset identification than ASSET when there are only significant marginal genetic effects. In addition to FUSION S2 and HUNT as identified by all pASTA methods, ASSET also includes D2D2007, METSIM, and TROMSO as associated studies, whose p-values with respect to marginal associations in study-specific analysis (Table S9) are far from significant. This data analysis demonstrates that without sacrificing much efficiency in the absence of G-E interaction, pASTA provides potential power gain when marginal genetic effects are modest and interactions are involved. Moreover, pASTA yields a more plausible subset of non-null studies.

### Application 2: North Finland Birth Cohort 1966 Data

We performed a genome-wide multiple-phenotype analysis by applying pASTA and ASSET to the lipid-related traits in the NFBC1966 study. Supplemental Table S10 presents the SNPs whose meta-analytic p-values are smaller than the genome-wide significance level (5×10^*-8*^). Seventeen SNPs are identified by pASTA(MA) at the genome-wide significance level. ASSET identifies seven SNPs as the genome-wide significant SNPs, and the only overlapping SNP between the two sets is rs754524, which is located in the *APOB* region on chromosome 2. However, it should be noted that since ASSET fails to produce p-values for some of the SNPs due to computational issues, the list of genome-wide significant SNPs generated by ASSET is not comprehensive and thus is not comparable to the list generated by pASTA(MA). Taking G-E interaction into consideration, pASTA(JOINT) identifies 13 SNPs, which are a subset of the SNPs identified by pASTA(MA). The SNP rs3764261, located on chromosome 16 near the *CETP* gene, is identified by pASTA(MA) (3.6×10^*-20*^) and pASTA(JOINT) (7.1×10^*-2O*^) as the top SNP. Figure 3 presents the QQ plots corresponding to the three methods.

We then compared the 17 genome-wide significant SNPs identified by pASTA(MA) to the 23 SNPs identified by another GWAS where all ten traits were analyzed jointly using a multivariate marginal association test (i.e. GEMMA) (Table S10). Seven SNPs belong to both lists, and the top five SNPs identified respectively by GEMMA and pASTA(MA) are the same (though in slightly different order).

We considered the top 100 SNPs identified by GEMMA for follow-up interaction analysis. The p-values produced by GEMMA for the 100 SNPs range from 5.5×10^*-30*^ to 7.0×10^*-5*^. The results of the combined p-values for the complete set is presented in Supplemental Table S11. Two SNPs, rs2083637 (on chromosome 8, near gene *LPL*) and rs754524 (on chromosome 2, in the *APOB* region), are used to illustrate the performance of the methods in the presence (rs2083637) or absence (rs754524) of G-E interaction. These two examples offer insight into the properties of the proposed methods and illustrate the advantage of incorporating G-E interaction into the analytic framework very clearly.

Supplemental Table S12 summarizes the linear regression coefficients and their corresponding 95% CI and p-values obtained from *Model 5* and *Model 6*. The SNP rs2083637, located near the *LPL* gene, is marginally associated with HDL and TG, with coefficients being −0.1 (*p* = 3.4×10^−6^*>*) and 0.1 (*p* = 5.6×10^−*6*^), respectively. CRP (*p* = 1.3×10^*-3*^), TC (*p* = 6.3×10^−*3*^), and LDL (*p* = 8.2×10^−3^) are associated with rs2083637 through SNP-BMI interactions. On the other hand, rs754524 has a positive marginal effect on HDL (*p* = 3.6×10^−2^) and a negative marginal effect on TC (*p* = 6.3×10^−10^), LDL (*p* = 2.1×10^−11^*>*), and TG (*p* = 1.8×10^−2^), while no significant SNP-BMI interactions are observed for these nine phenotypes under the 0.05 threshold.

For rs2083637, ASSET and pASTA(MA) both identify TG and HDL as associated phenotypes (Table 4). However, both pASTA(JOINT) and pASTA(MA+CC) identify three additional phenotypes (CRP, TC, LDL), for which rs2083637 has significant interaction with BMI, when G-E interaction is taken. into account. Supplemental Figure S5 presents the effect and the corresponding 95% CIs for rs2083637 on CRP, TC, and LDL when BMI equals the mean, mean±1 standard deviation (sd), and mean±2sd. We observe variation of the SNP effect sizes across different levels of BMI. For example, rs2083637 has positive but not significant effect on CRP for subjects with average BMI, while for subjects whose BMI is 2 sd greater than the average, the association becomes negative and significant at the level of 0.05. Therefore, the incorporation of G-E interaction can enhance the knowledge that is not available in marginal association testing. The overall p-value of associations given by pASTA(JOINT) (2.5×10^−6^) is also smaller than those given by 1-df methods (2.7×10^−5^ for pASTA[MA] and 3.6×; ^−6^for ASSET). For rs754524, pASTA(MA) identifies two phenotypes (TC and LDL), both of which have significant marginal associations with this SNP in phenotype-specific analysis, as associated phenotypes. SNP rs754524 has been previously reported to be associated with both traits. [31,32]. ASSET identifies two additional phenotypes, glucose and HDL, and fails to identify TC. The evidence for the marginal association of this SNP with HDL is not strong [27], which indicates that such identified association may be false. pASTA (JOINT) identifies the same set of associated traits as pASTA(MA). This example shows that by incorporating interactions one can obtain smaller p-values for detecting overall associations and discover more relevant traits that are missed by ASSET, ones that show significant G-E interaction effects in phenotype-specific analysis.

**Table 4.**
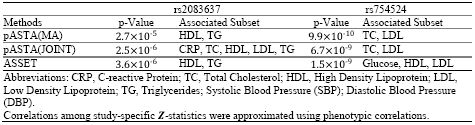
Joint Analysis of Multiple Phenotypes in NFBC 1966 Study

## Discussion

We propose a powerful subset-based framework for the analysis of association studies after accounting for potential G-E interactions. Simulation studies demonstrate that this framework improves the power to detect overall associations as well as the accuracy of determining associated studies/traits by incorporating G-E interactions, while maintaining comparable power to ASSET in the absence of such interactions. Our data examples exemplify that 2-df pASTA yields smaller meta-analytic p-values for SNPs that show significant G-E interactions in study/phenotype-specific analysis compared to 1-df methods where only marginal genetic associations are considered (Table 3 and Table 4). In addition, our analysis on the NFBC1966 data illustrates that pASTA is able to identify new phenotypes that are associated with given susceptibility loci after considering G-E interactions that will be missed by traditional marginal association analysis (Table 4). These properties make pASTA a powerful tool to identify candidate SNPs that are potentially associated with the trait(s) of interest in the screening step of a GWAS, which then can be followed by deeper characterization/interpretation of the associations through other methods, such as biological validation. For example, RNA sequencing experiments can be performed to examine whether the candidate SNPs affect the expression level of the target gene [33]. Crispr-Cas9 knockout screening can be carried out in human cell lines to determine the functional significance of these candidate SNPs [34].

Our simulation studies show that in the presence of interaction, pASTA(MA+EB) has the best performance in terms of all four evaluation metrics in a case-control setting where G-E independence holds. The rest of the 2-df methods (Fisher’s methods, pASTA[JOINT] and pASTA[MA+CC]) give similar power for detecting overall associations across the scenarios (Figures 1B-1E), but pASTA has the advantage of being able to identify the subset of studies/traits that are most likely to contribute to the associations. For a cohort study with multiple phenotypes where the G-E independence does not apply, pASTA(JOINT) gives the best performance in terms of the four metrics (Figures 2E-2H).

There are several limitations of our proposed approach that lend the problem to future research. The present proposal for pASTA focuses on testing the existence of genetic associations using p-values as inputs and is purely bi-directional in nature. It aggregates the evidence of both genetic effects and G-E interactions from individual studies but does not specify the directions of such effects or interactions. It is a pure testing approach and does not provide pooled regression coefficient estimates from meta-analysis. Thus it loses insight into the nature of the environmental heterogeneity across cohorts. Further work is needed to define the subsets of studies that have similar direction of genetic effects in subsets defined by the environmental factor (for example the sign of estimates *β*_*G*_ in the group *E* = 0 and of *β*_*GE*_+*β*_*G*_ in the group *E* = 1 for a binary exposure E). The method is currently based on the assumption of fixed effect(s) meta-analysis. Since the input is p-values, the method can be extended to random effects meta-analysis as well as hybrid methods that combine fixed and random effects meta-analysis [35]. In our simulation studies and data analysis, pASTA uses phenotypic correlations to approximate the correlations among *Z*_*k*_’s, which may result in compromised power and conservative type I error. Optimizing the estimation of such correlation is an issue that needs to be further addressed.

We only focused on candidate loci with known marginal association or gene-environment interaction in our data examples. Simulation results indicate that pASTA is scalable to a genome-wide level and maintains Type 1 error at smaller 10^*-*5^ threshold of α. In the T2D data analysis, the average run time of pASTA for the marginal effect, i.e. pASTA(MA), for one SNP was 181.7 milliseconds on a 2.9 GHz Intel processor, and the incorporation of G-E interaction, i.e. pASTA(JOINT), yielded a similar computation time (159.2 milliseconds). For the data from the NFBC 1966 study where correlations across the test statistics need to be taken into account, the average run times for both pASTA(MA) and pASTA(JOINT) for one SNP were 1.7 seconds. For a typical GWAS to analyze one million SNPs, the projected computation time of pASTA on 20 CPU cores is approximately one day. ASSET was slightly faster than pASTA in the NFBC 1966 data analysis (1.5 seconds) but twice slower in the T2D data analysis (3.3 seconds). Computing time was calculated using the R package microbenchmark [36]. We plan to optimize the pASTA implementation code and develop an efficient R package akin to ASSET.

## Appendix

### Detailed Derivation of Meta-analytic p-value

For an observed test statistic *Z*_*meta*_ *= T*_*obs*_, the DLM approximated p-value can be expressed as

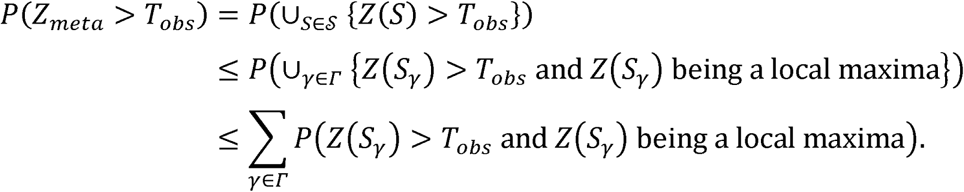

For a fixed subset *S*_*γ*_ ⊆ {1,…, *K*},

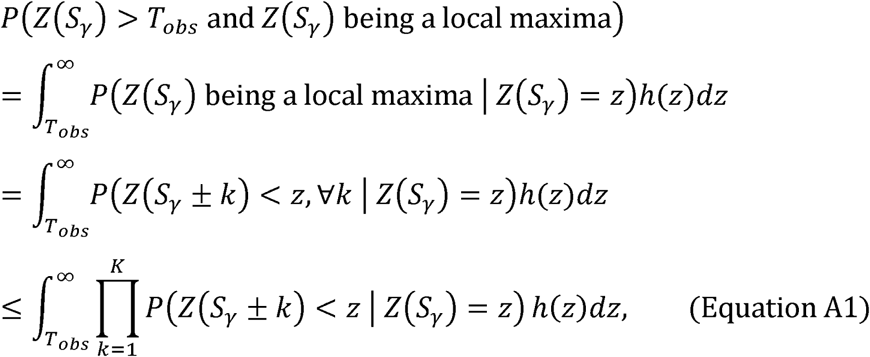

where h(·) is the probability distribution function of *Z*(*S*_*γ*_) The last inequality is justified by the “separability” assumption that given the current subset, its neighbors have independent *Z*-statistics.

i. When *k* ∉ *S*_*γ*_, given that Z(S_*γ*_) = *z*,

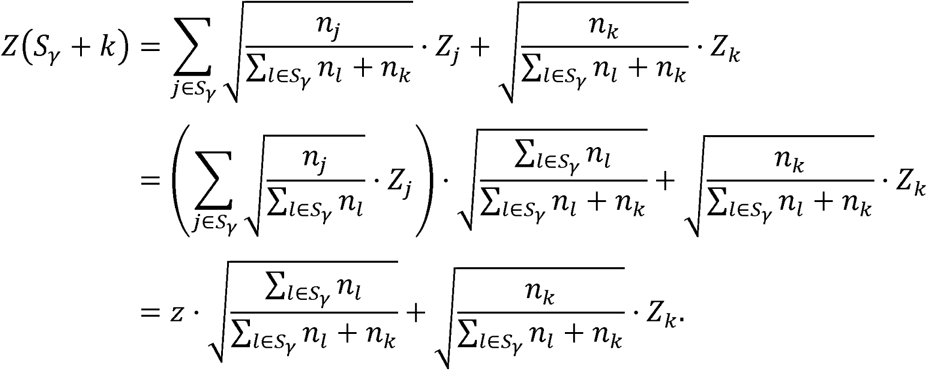 Hence,

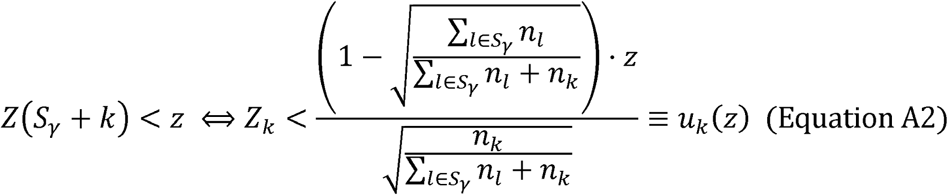

and

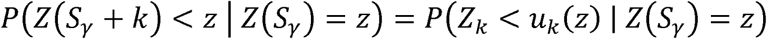
ii. When *k* ∈ *S*_*γ*_, given that Z(S_*γ*_) = *z,*

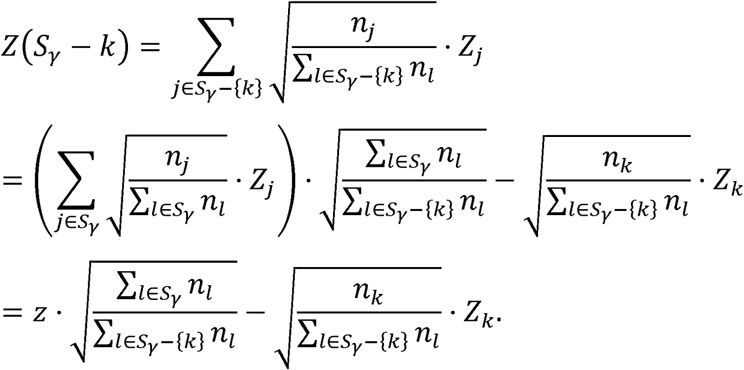 Hence,

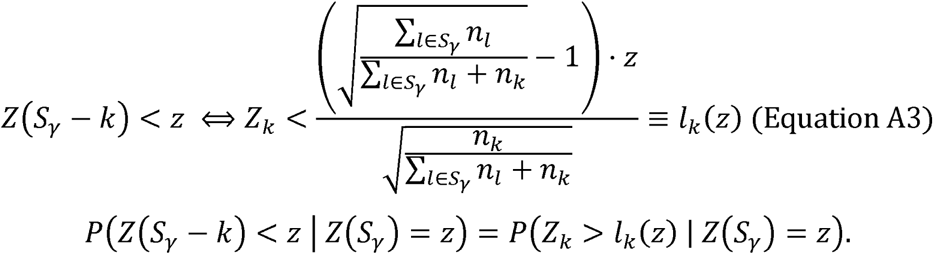 This conditional probability of *Z*_*k*_ given *Z*(*S*_*γ*_) can be evaluated based on a joint bivariate normal distribution, which is specified in the main text. For numerical evaluation of the integrals, we directly apply the R function integrate().

### Metrics Used in Power Analysis

We used two main metrics for evaluation:

- **Power of detecting genetic associations that truly exist,** defined as the probability of rejecting the relevant null hypotheses *H*_ok_ within study *k* with a pre-specified level *α*. This quantity is estimated as the empirical proportion of replications where the meta-analytic p-value combined across all studies is smaller than *α*
- **Power of identifying the exact true subset of non-null studies,** defined as the probability of correctly identifying such subset. The definition of non-null studies is specified above. The corresponding empirical proportion of replications declaring the correct subset to be non-null is used to estimate this power.

We also report the sensitivity and specificity of identifying the subset of non-null studies, defined as the proportion of non-null studies that are correctly identified by subset-based approaches, and the proportion of truly null studies that are declared null, respectively. They are both estimated by the corresponding empirical proportions in the simulation. Note that sensitivity and specificity respectively allow false positives and false negatives in the identified subset, while power of identifying exactly the true subset is a much more stringent criterion.

## Acknowledgement

The research of BM was supported by the National Science Foundation grant DMS 1406712, the research of SL was supported by the National Institute of Health grant R01-HG008773, and the research of MB was supported by HG009976. The FUSION study was supported by DK062370. We thank the D2D2007, DIAGEN, FUSION S2, HUNT, METSIM, and TROMSO investigators for providing access to their data. The NFBC1966 Study is conducted and supported by the National Heart, Lung, and Blood Institute (NHLBI) in collaboration with the Broad Institute, UCLA, University of Oulu, and the National Institute for Health and Welfare in Finland. This manuscript was not prepared in collaboration with investigators of the NFBC1966 Study and does not necessarily reflect the opinions or views of the NFBC1966 Study Investigators, Broad Institute, UCLA, University of Oulu, National Institute for Health and Welfare in Finland and the NHLBI.

The authors declare no conflicts of interest.

## Web Resources

Current annotated code, https://github.com/youfeiyu/pASTA/

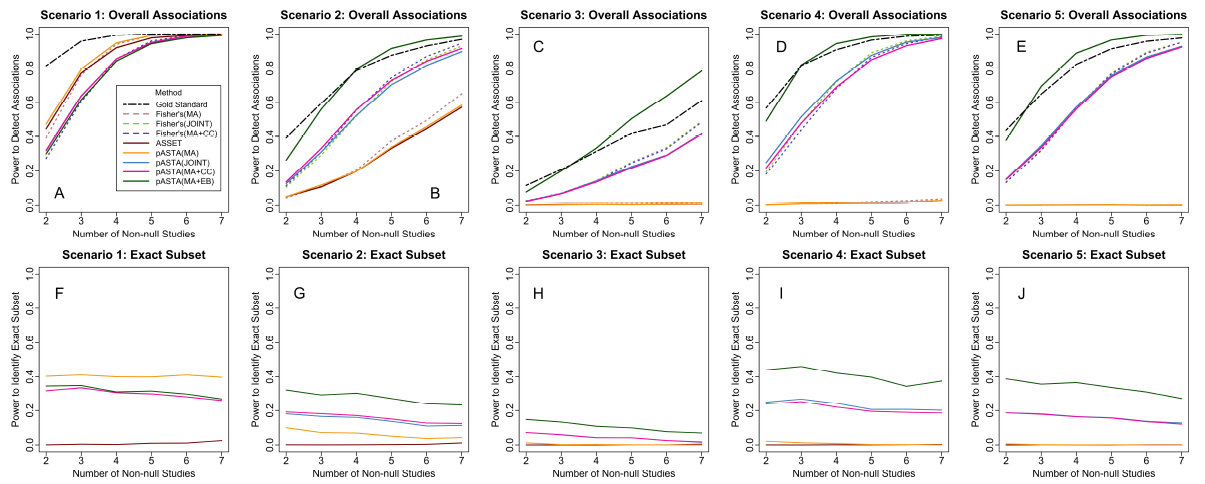

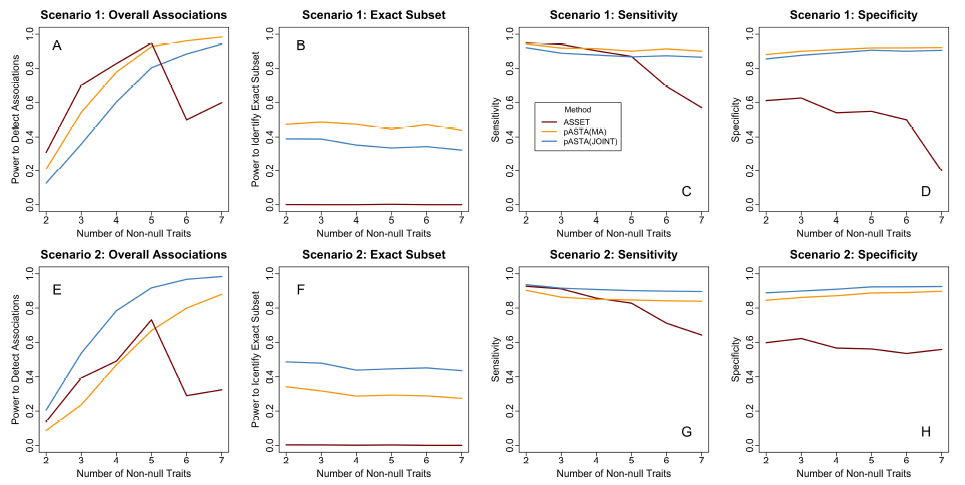

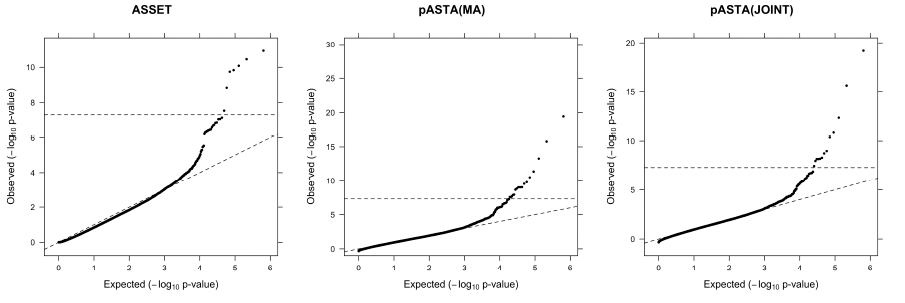

